# Human iPSC-derived alveolar and airway epithelial cells can be cultured at air-liquid interface and express SARS-CoV-2 host factors

**DOI:** 10.1101/2020.06.03.132639

**Authors:** Kristine M. Abo, Liang Ma, Taylor Matte, Jessie Huang, Konstantinos D. Alysandratos, Rhiannon B. Werder, Aditya Mithal, Mary Lou Beermann, Jonathan Lindstrom-Vautrin, Gustavo Mostoslavsky, Laertis Ikonomou, Darrell N. Kotton, Finn Hawkins, Andrew Wilson, Carlos Villacorta-Martin

## Abstract

Development of an anti-SARS-CoV-2 therapeutic is hindered by the lack of physiologically relevant model systems that can recapitulate host-viral interactions in human cell types, specifically the epithelium of the lung. Here, we compare induced pluripotent stem cell (iPSC)-derived alveolar and airway epithelial cells to primary lung epithelial cell controls, focusing on expression levels of genes relevant for COVID-19 disease modeling. iPSC-derived alveolar epithelial type II-like cells (iAT2s) and iPSC-derived airway epithelial lineages express key transcripts associated with lung identity in the majority of cells produced in culture. They express *ACE2* and *TMPRSS2*, transcripts encoding essential host factors required for SARS-CoV-2 infection, in a minor subset of each cell sub-lineage, similar to frequencies observed in primary cells. In order to prepare human culture systems that are amenable to modeling viral infection of both the proximal and distal lung epithelium, we adapt iPSC-derived alveolar and airway epithelial cells to two-dimensional air-liquid interface cultures. These engineered human lung cell systems represent sharable, physiologically relevant platforms for SARS-CoV-2 infection modeling and may therefore expedite the development of an effective pharmacologic intervention for COVID-19.

## Introduction

In December 2019, a previously unknown infectious illness emerged in Wuhan, Hubei Province, China (Zhu 2020), with the most severe cases progressing to acute respiratory distress syndrome (ARDS), cytokine storm syndrome, and death (Huang 2020, Mehta 2020). A novel coronavirus, Severe Acute Respiratory Syndrome (SARS)-coronavirus-2 (SARS-CoV-2) was identified as the pathogen that causes the illness now termed “Coronavirus Disease 2019” (COVID-19) (Zhu 2020). COVID-19 rapidly spread through the population via human-to-human transmission (Chan 2020, Zhu 2020). The World Health Organization declared COVID-19 a Public Health Emergency of International Concern on January 30, 2020, and on March 11 characterized the outbreak as a pandemic. By May 28, 2020, 216 countries, areas, or territories around the world have cumulatively reported 5.6 million cases, with over 350,000 deaths (WHO Coronavirus Disease Dashboard).

Examination of the lungs of fatal cases of COVID-19 using transmission electron microscopy identified SARS-CoV-2 viral particles in alveolar epithelial type 2 cells (AT2s), type 1 cells, and epithelial cells of the trachea (Bradley 2020). Lung histology was characterized by typical ARDS findings including diffuse alveolar damage and evidence of airway inflammation in COVID-19 autopsy studies (Bradley 2020, Barton 2020, Zhang Allergy 2020). Outside of the lung, there is increasing evidence that gastrointestinal involvement in SARS-CoV-2 infection may also contribute to both disease severity and transmission (Wang 2020, Xu 2020).

In the face of this pandemic and despite intense COVID-19-focused research efforts, there is a pressing need for physiologically relevant human model systems that can be used to study the pathogenesis of SARS-CoV-2 infection and develop therapeutic strategies. Given the surge in COVID-19 research, such a model would also be scalable to meet the demands of the research community.

Pre-clinical models of pulmonary viral infection and drug screening platforms typically rely on human lung cancer-derived cell lines. For example, Calu-3 cells, originally derived from a human lung adenocarcinoma, were used as a respiratory epithelial surrogate to model coronavirus infections including SARS (Tseng 2005, Chan 2013), Middle East Respiratory Syndrome (MERS) (Lau 2013, Chu 2018), and SARS-CoV-2 (Hoffmann 2020, Zhang Science 2020). However, the genomes of Calu-3 and other common lung epithelial cell lines are known to have accumulated numerous aberrations, have drifted away from displaying a pulmonary epithelial molecular phenotype (Klijn 2015), and therefore are unlikely to faithfully model the pulmonary epithelial response to SARS-CoV-2 infection.

Induced pluripotent stem cell (iPSC) technology offers the potential to provide human cells more reminiscent of human airway and alveolar epithelial cells for SARS-CoV-2 research. Our group has published directed differentiation protocols for generating iPSC-derived lineages of interest in SARS-CoV-2 infection, including lung airway (Hawkins 2020), lung alveolar (Jacob 2017), and intestinal (Mithal 2020) epithelial cells. iPSC-derived cells generated from noninvasively obtained patient samples, such as peripheral blood, can be banked and used to generate multiple organ lineages from a single individual. iPSC-derived lung and intestinal epithelial cells derived in our protocols are transcriptomically similar to their respective primary counterparts, recapitulate appropriate cell-intrinsic phenotypes of genetic disease, and can respond to immune stimuli (Hawkins 2020, Jacob 2017, Mithal 2020). iPSC-based systems provided key insights into the cellular-level pathological mechanisms underlying diseases and generated novel drug candidates, some of which have progressed to clinical trials (Shi 2017).

Here, we define the gene expression profiles of iPSC-derived epithelial cell types focusing on those transcripts that encode genes considered essential for modeling SARS-CoV-2 infection. First, we compare current lung cancer cell lines used to study respiratory pathogens to primary and iPSC-derived lung epithelial cells and determine that cancer lines lack expression of several fundamental lung epithelial genes. We establish a novel iPSC-derived alveolar epithelial type 2 cell (iAT2) air-liquid interface (ALI) culture system to enable modeling of environmental exposures of the human alveolar epithelium, including viral infection. Importantly, we observe the expression of *ACE2* and *TMPRSS2*, two genes encoding host cell proteins essential for SARS-CoV-2 cell entry (Hoffmann 2020) in iPSC-derived airway, alveolar, and intestinal epithelial cells.

## Results

### Human lung cancer cell lines do not recapitulate the transcriptomic program of primary AT2s

Human lung cancer-derived cell lines are commonly used as pre-clinical models of the lung epithelium. To understand the suitability of lung cancer cell lines compared to iAT2s as an *in vitro* model of the alveolar epithelium, we evaluated the extent to which cancer cell lines and iAT2s transcriptomically resemble primary AT2s. We compared expression of key AT2 genes in RNA sequencing data from primary human adult AT2s and iAT2s (Jacob 2017, Hurley 2020) to RNA sequencing data from 150 lung cancer cell lines (Klijn 2015). The cell lines Calu-3, A549 and H1299, described in SARS-CoV-2 infection studies, were included (Hoffmann 2020, Letko 2020, Matsuyama 2020, Zhang Science 2020). We found that Calu-3, A549, and H1299 cells do not express detectable *NKX2-1* transcript, a key lung epithelial transcription factor that is expressed by adult AT2s and iAT2s (Fig 1) as well as adult airway basal and airway secretory cells (Hawkins 2020, McCauley 2017), and is required for the expression of many lung epithelial-specific transcripts. No cell lines expressed all of the AT2 transcriptomic program components *SFTPC, SFTPD, SFTPA1, SFTPA2*, and *PGC*, while all were expressed in adult AT2s and iAT2s (Fig 1).

**Figure 1.**
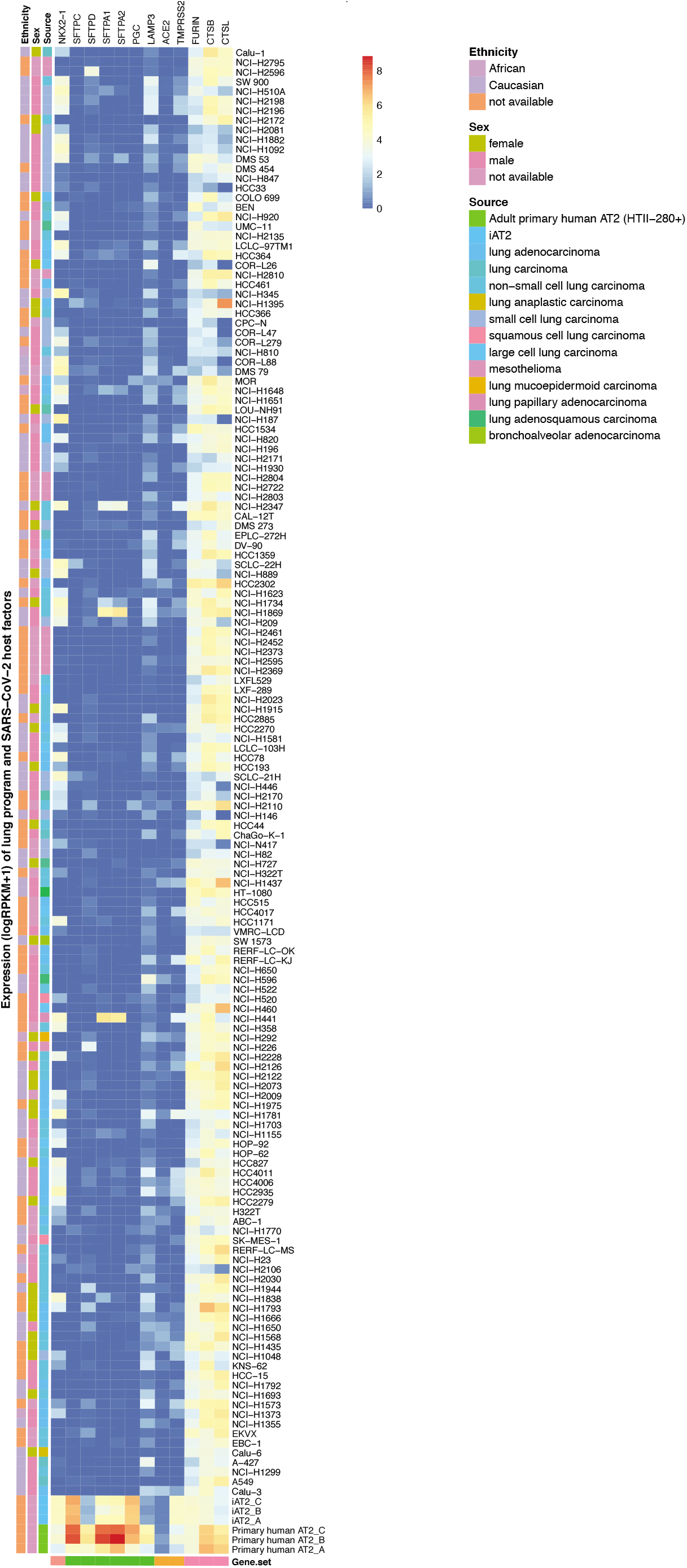
Human lung cancer cell lines do not recapitulate the transcriptomic program of primary AT2s. Heatmap with the expression of AT2 markers genes and SARS-CoV-2 host factors in primary human adult AT2 cells, iAT2 cells, and lung cell lines commonly used for pre-clinical models (ArrayExpress E-MTAB-2706).

### iAT2s maintain AT2-like identity at air-liquid interface

Primary human bronchial epithelial cells, consisting predominantly of airway basal cells, can be serially passaged in 3D or 2D cultures and have been extensively characterized after differentiation into a variety of airway lineages through culturing in a 2D ALI. Similarly, we have demonstrated that human iPSC-derived airway basal cells (iBCs) can be maintained and passaged in 3D culture or differentiated in 2D ALI cultures (Fig 2) (Hawkins 2020). iBCs express airway basal cell markers TP63/NGFR/KRT5 and give rise to other airway lineages such as secretory cells (SCGB1A1+; MUC5AC+) and ciliated cells (FOXJ1+; ACT+) when expanded in differentiation media in 3D or plated in ALI culture (Hawkins 2020). iPSC-derived airway cells recapitulate airway functionality and model disease-relevant phenotypes such as CFTR-mediated chloride transport and mucus cell metaplasia upon IL-13 exposure (Hawkins 2020).

**Figure 2.**
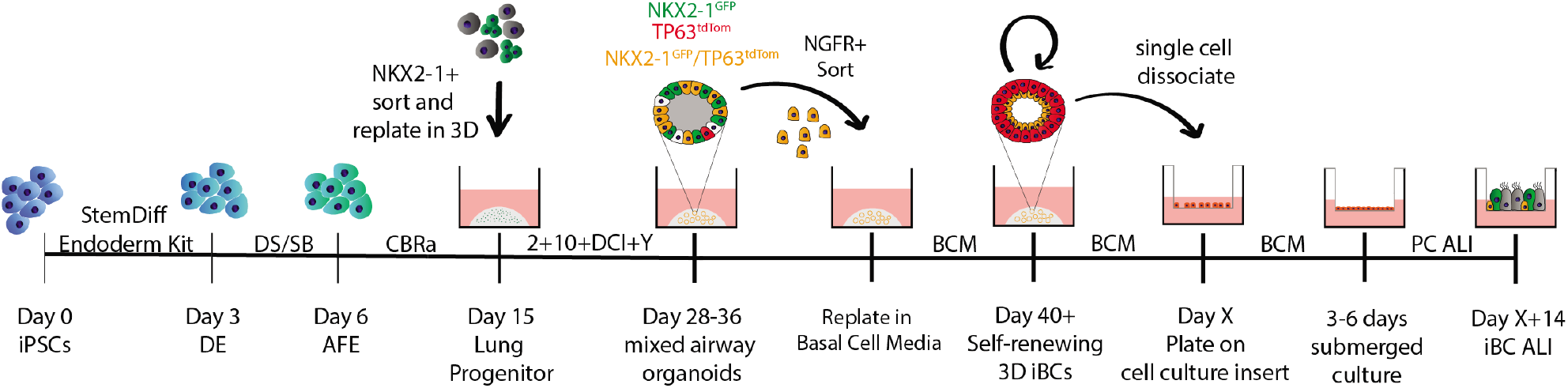
iBCs can be maintained in 3D culture and differentiated at ALI. Schematic displaying directed airway differentiation into ALI as outlined in Hawkins et al., BioRxiv 2020. DE = Definitive Endoderm, AFE = Anterior Foregut Endoderm. DS/SB = 2uM dorsomorphin + 10uM SB431542, CBRa = 3uM CHIR99021 + 10 ng/ml BMP4 + 100nM retinoic acid, DCI = 50nM dexamethasone + 0.1mM cAMP + 0.1mM IBMX, Y = 10uM Rho-associated kinase inhibitor (Y-27632). BCM = Basal Cell Media comprised of Pneu-maCult-Ex Plus medium supplemented with the SMAD inhibitors. A831 and DMH1, and Y-27632.

Although ALI cultures of primary airway epithelial cells have been used to successfully model viral infections (Escaffre 2016, Griggs 2017, Persson 2014), primary human AT2 cell cultures are difficult to maintain in self-renewing cultures, currently require mesenchymal feeders for passaging, and thus have not been routinely employed for viral infection models. We sought to adapt our previously published human iAT2 culture system which involved the culture of iAT2s as monolayered epithelial spheres suspended in Matrigel to ALI culture. iAT2s were plated on Transwell filters in serum-free culture medium without feeders and formed an intact epithelial layer allowing removal of the medium from the apical chamber and establishment of ALI (Fig 3A).

**Figure 3A-C.**
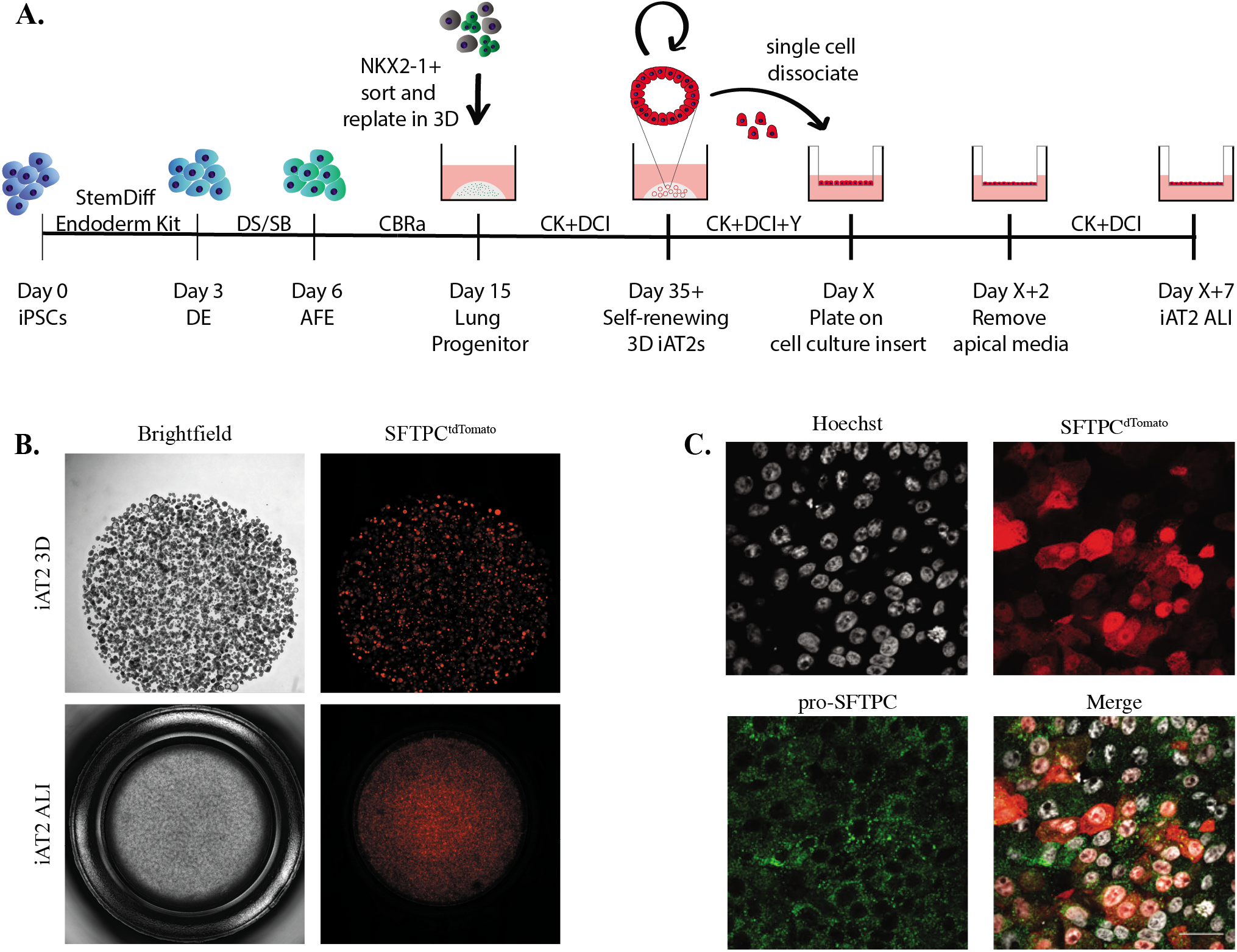
iAT2s maintain AT2-like identity when cultured at air-liquid interface. (A) Differentiation schematic to generate 3D iAT2s and plate at ALI. iAT2s cultured as 3D spheres were dissociated and re-plated on cell culture inserts to generate ALI cultures. Apical media was removed after 2 days in submerged culture. ALI cultures could be maintained for 14-21 days post passage. DE = Definitive Endoderm, AFE = Anterior Foregut Endoderm. DS/SB = 2uM dorsomor-phin + 10uM SB431542, CBRa = 3uM CHIR99021 + 10 ng/ml BMP4 + 100nM retinoic acid, CK = 3uM CHIR99021 + 10ng/ml KGF, DCI = 50nM dexamethasone + 0.1mM cAMP + 0.1mM IBMX, Y = 10uM Rho-associated kinase inhibitor (Y-27632). (B) iAT2s generated from iPSCs targeted with a tdTomato-encoding cassette to the endogenous SFTPC locus express tdTomato when cultured in 3D spheroids (iAT2 3D) and at ALI (iAT2 ALI). (C) iAT2s cultured at ALI express the SFTPC^tdTomato^ reporter as well as pro-SFTPC by immunofluo-rescent staining.

After establishing this ALI protocol, we sought to understand whether iAT2s maintain AT2-like identity at ALI. iAT2s were generated from an iPSC line (SPC2; Hurley 2020) carrying a tdTomato-encoding cassette targeted to the *SFTPC* locus. These iAT2s expressed SFTPC^tdTomato^ in 3D spheroid culture as well as at ALI (Fig 3B). Similarly, iAT2s at ALI expressed pro-SFTPC protein based on immunofluorescent staining (Fig. 3C). The majority of iAT2s cultured at ALI expressed the key lung epithelial transcription factor NKX2-1 by flow cytometry analysis (94% ± 1.5%) (Fig. 3D).

**Figure 3D-F.**
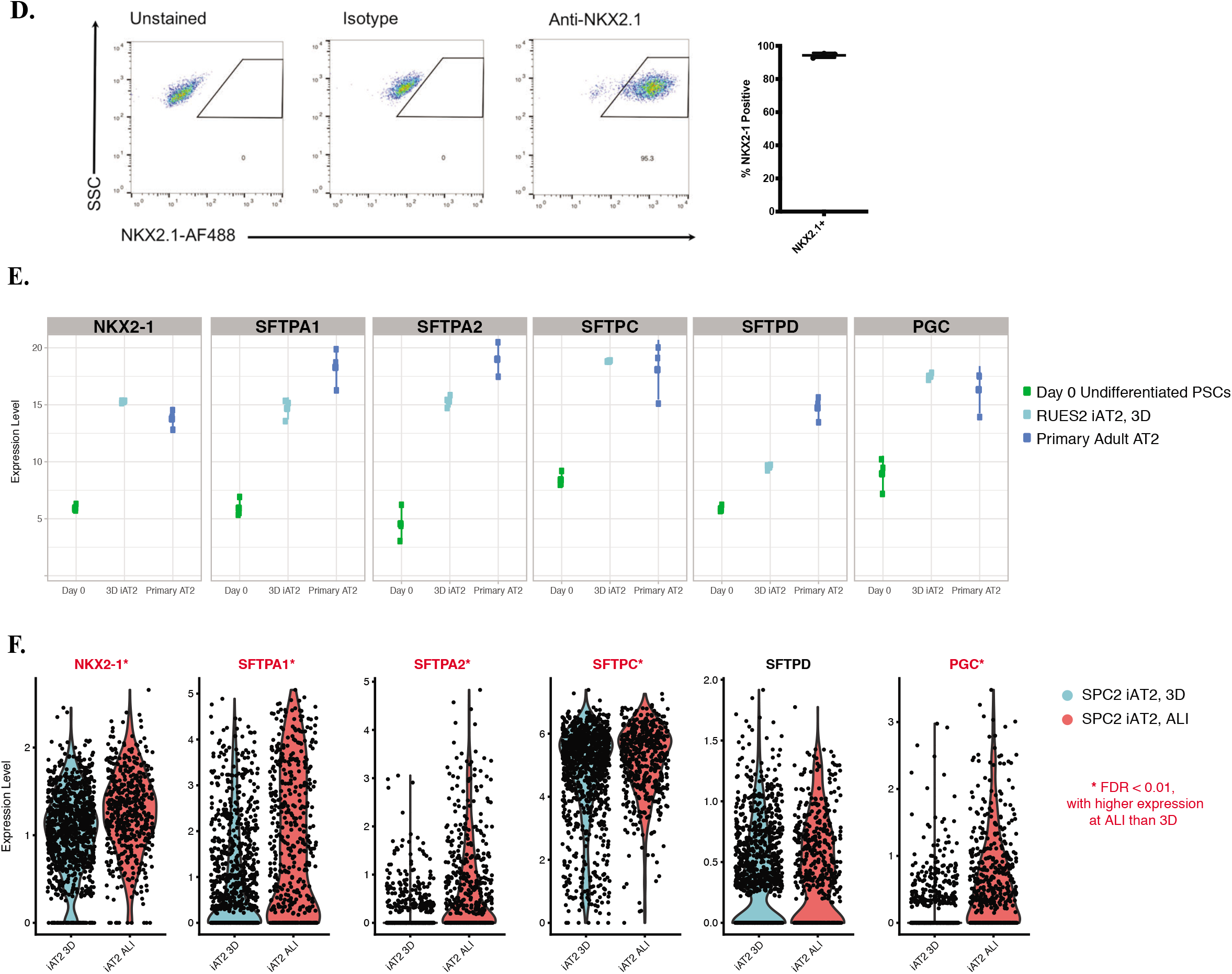
iAT2s maintain AT2-like identity when cultured at air-liquid interface. (D) 94% +/- 1.5% of iAT2s cultured at ALI are NKX2-1+ by flow cytometry analysis. (E) Day 0 (undifferentiated RUES2 PSCs), RUES2 iAT2s cultured as 3D spheres, and primary adult HTII-280-sorted AT2s were profiled by RNA-Seq head-to-head. iAT2s cultured in 3D spheres express *NKX2-1, SFTPC*, and *PGC* at levels similar to primary adult AT2s. iAT2s express *SFTPA1, SFTPA2*, and *SFTPD* at slightly lower levels than primary adult AT2s. (F) iAT2s generated from the SPC2 iPSC line were cultured in 3D or ALI in parallel and profiled by scRNA-Seq. iAT2s cultured at ALI express *NKX2-1, SFTPC*, and *SFTPD* at similar levels to iAT2s cultured in 3D spheres. iAT2s at ALI express significantly increased levels of *NKX2-1, SFTPA1, SFTPA2, SFTPC*, and *PGC* compared to 3D (FDR < 0.01), and similar levels of *SFTPD*.

We have previously compared the transcriptomes of RUES2-derived iAT2s cultured as 3D spheres to undifferentiated human pluripotent stem cells (hPSCs, “RUES2 day 0”) and adult HTII-280+ primary AT2 controls (n=3 separate lungs; Jacob 2017, Hurley 2020). iAT2s cultured as 3D spheres express the key lung transcription factor *NKX2-1* and AT2 consensus signature genes *SFTPA1, SFTPA2, SFTPC, SFTPD*, and *PGC* (Fig. 3E). We performed single-cell RNA sequencing (scRNA-Seq) of SPC2-derived iAT2s cultured in parallel as either 3D spheres or at ALI, and found that iAT2s at ALI express all 6 of these marker genes at similar or significantly higher levels compared to 3D culture (Fig. 3F).

### iAT2s and iBC-derived airway epithelial cells are transcriptomically similar to primary counterparts

In order to explore the overall similarity between samples, we integrated iPSC-derived airway and alveolar epithelial cells with their primary counterparts. We merged the scRNA-Seq dataset from iAT2s cultured at ALI and 3D (Fig. 3F) with two previously published scRNA-Seq datasets; one of iBCs cultured at ALI (Hawkins 2020) and another of human adult lung epithelial cells (Habermann 2019) (Fig. S1).

iAT2s cultured in 3D and at ALI cluster separately from other cell types, and iAT2s at ALI overall cluster closest to primary AT2s compared to other adult cell types (Fig. 4A). iBCs cultured at ALI differentiate to secretory and ciliated-like cells (Hawkins 2020); each iPSC-derived airway epithelial cell type represented in the culture system (iBasal, iSecretory, and iCiliated) clusters with the corresponding primary cell type (basal, secretory, and ciliated) (Fig. 4A) (Hawkins 2020).

**Figure 4.**
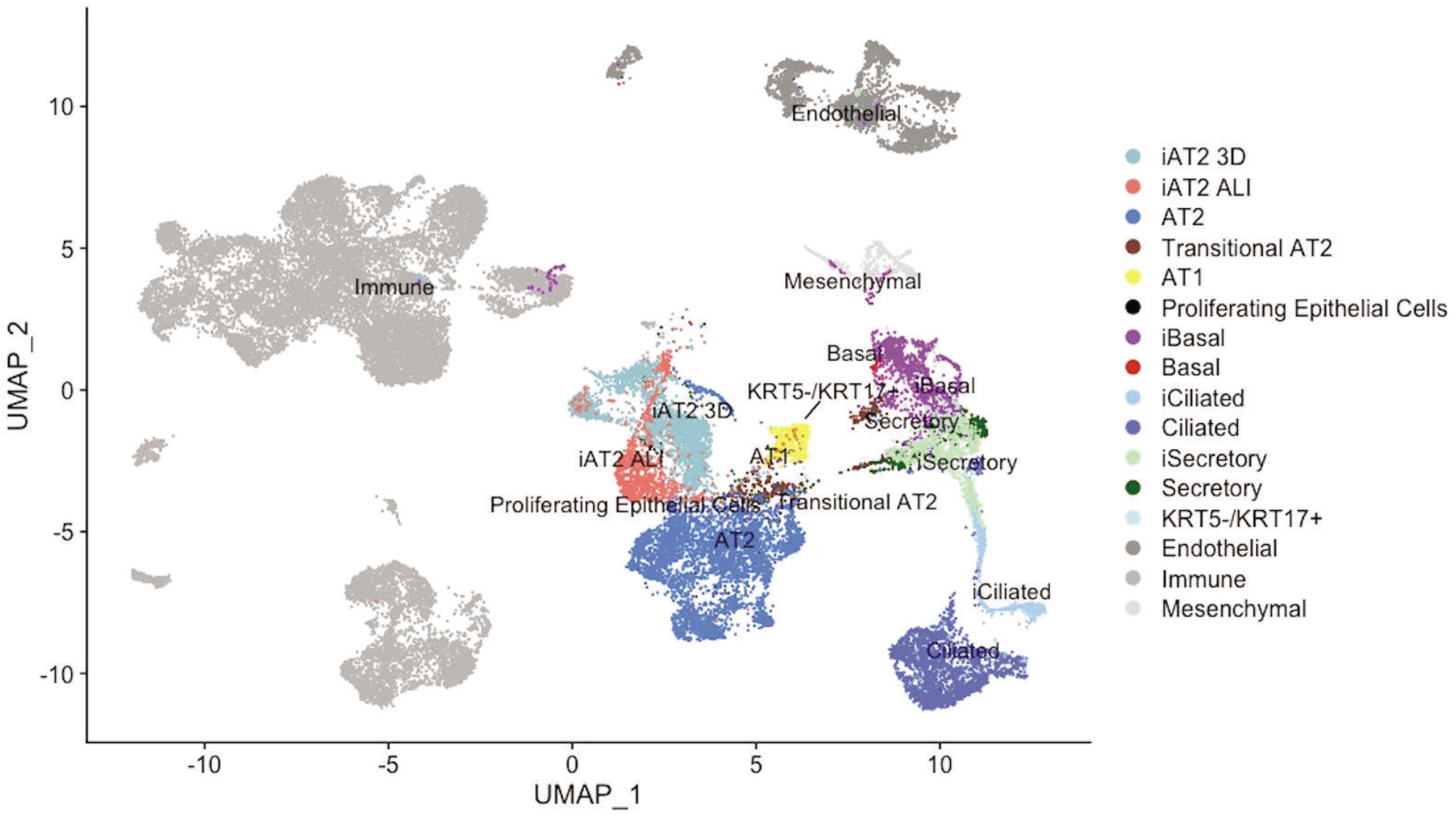
iAT2s and iBC-derived airway epithelial cells are transcriptomically similar to primary counterparts. UMAP of integrated analysis of an adult human lung dataset (Habermann 2019), iBC ALI cultures (Hawkins 2020), and iAT2s cultured at ALI and in 3D.

### iAT2s and iBC-derived airway epithelial cells express *ACE2* and *TMPRSS2*

Expression levels of the putative host factors required for SARS-CoV-2 cellular entry, ACE2 and TMPRSS2, are potential indicators of cell susceptibility to viral infection (Hoffman 2020) and therefore model suitability. To assess whether iAT2s express *ACE2* and *TMPRSS2*, we compared RUES2-derived iAT2s cultured as 3D epithelial spheres to freshly sorted adult HTII-280+ primary AT2s (n=3 separate adult lungs) that were profiled head to head in our previously published RNA-Seq dataset without the potentially confounding batch effects that may be introduced when merging datasets procured in separate experiments (Jacob 2017, Hurley 2020). We observed that 3D iAT2s and primary human adult AT2s express similar levels of both *ACE2* and *TMPRSS2* (Fig 5A).

**Figure 5.**
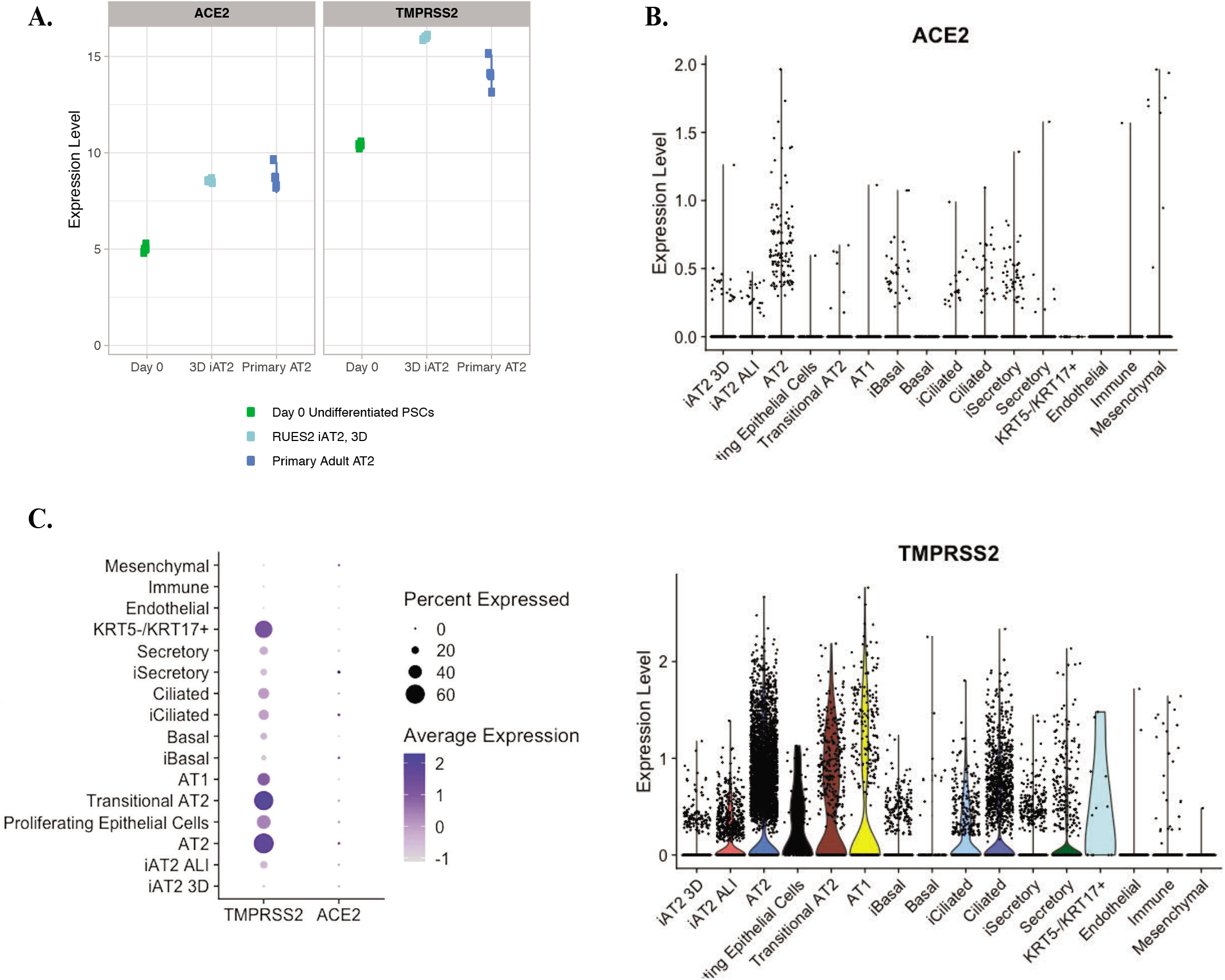
iAT2s and iBC-derived airway epithelial cells express *ACE2* and *TMPRSS2*. (A) Day 0 (undifferentiated RUES2 PSCs), RUES2 iAT2s cultured as 3D spheres, and primary adult HTII-280-sorted AT2s were profiled by RNA-Seq. iAT2s cultured as 3D spheres express ACE2 and TMPRSS2 at similar levels compared to primary adult AT2s. (B) Level of expression of ACE2 and TMPRSS2 in iAT2s cultured in 3D (iAT2 3D) and at ALI (iAT2 ALI), iBC-derived cells (iBasal, iCiliated, and iSecretory), and primary human lung cells (AT2, Proliferating Epithelial Cells, Transitional AT2, AT1, Basal, Ciliated, Secretory, KRT5-/KRT17+, Endothelial, Immune, Mesenchymal). (C) Percentage of cells expressing ACE2 and TMPRSS2 in iAT2s cultured in 3D (iAT2 3D) and at ALI (iAT2 ALI), iBC-derived cells (iBasal, iCiliated, and iSecretory), and primary human lung cells (AT2, Proliferating Epithelial Cells, Transitional AT2, AT1, Basal, Ciliated, Secretory, KRT5-/KRT17+, Endothelial, Immune, Mesenchymal).

Next, we examined the scRNA-Seq datasets described previously to determine the level and relative proportion of *ACE2* and *TMPRSS2*-expressing cells in iAT2s cultured in 3D, iAT2s cultured at ALI, iBC-derived ALI cultures, and primary human lung epithelial cells. We observed *ACE2* and *TMPRSS2* expression in a wide variety of lung epithelial lineages, including primary AT2s, iAT2s, iAT2s after ALI culture, iBCs, iCiliated cells, and iSecretory cells (Fig 5B-C). Cells that express *ACE2* mRNA were rare in both primary and iPSC-derived datasets: 2.6% of primary AT2s, 1% of iAT2s cultured in 3D, and 1.1% of iAT2s cultured at ALI (Fig 5B-C). This frequency of expression is similar to the previously reported 1.4% *ACE2+* AT2s in human lung scRNA-Seq data (Ziegler 2020). In contrast, 62.2% of primary AT2s express *TMPRSS2* transcript compared to 3.3% of iAT2s cultured in 3D and 19.7% of iAT2s cultured at ALI (Fig 5B-C). Others have reported a frequency of 34.2% *TMPRSS2+* AT2s in human lung scRNA-Seq data (Ziegler 2020).

We compared the expression of *ACE2* and *TMPRSS2* in iPSC-derived airway cells to expression in those distal airway lineages captured by Habermann et al. Although the primary cells captured from this distal lung dataset are relatively depleted in large airway secretory, ciliated, and basal cells, *ACE2* was expressed in 0%, 0.5%, and 1% of these *distal* primary basal, secretory, and ciliated cells, compared to expression in 2.6%, 4.2%, and 3.1% of iBC, iSecretory, and iCiliated cells, respectively (Fig 5B-C). iPSC-derived basal, secretory, and ciliated cells expressed *TMPRSS2* at proportions of 11.4%, 16.7%, and 29.1% compared to primary cell counterparts which expressed at proportions of 16.9%, 22.6%, and 32.0% (Fig 5B-C).

### iPSC-derived Human Intestinal Organoids (HIO) express *ACE2* and *TMPRSS2*

Extrapulmonary cell types such as intestinal epithelial cells have been indicated to be susceptible to SARS-CoV-2 infection (Zhang Allergy 2020, Holshue 2020). Therefore, we sought to test whether our previously described iPSC-derived HIO system (Mithal 2020) expresses the key SARS-CoV-2 host factor genes *ACE2* and *TMPRSS2*.

Differentiated HIOs (day 42 of differentiation) and their intestinal progenitor-like precursors (day 8 of differentiation) were profiled by digital gene expression (Mithal 2020).

Differentiated HIOs significantly upregulate both *ACE2* and *TMPRSS2* compared to day 8 intestinal progenitor-like cells (Fig. 6A). To contextualize this finding, we subsequently performed qRT-PCR for *ACE2* and *TMPRSS2* in undifferentiated iPSCs, differentiated HIOs, and a primary human adult colon sample. Preliminary results suggest HIOs express *ACE2* and *TMPRSS2* to a level comparable to primary human colon (Fig. 6B).

**Figure 6.**
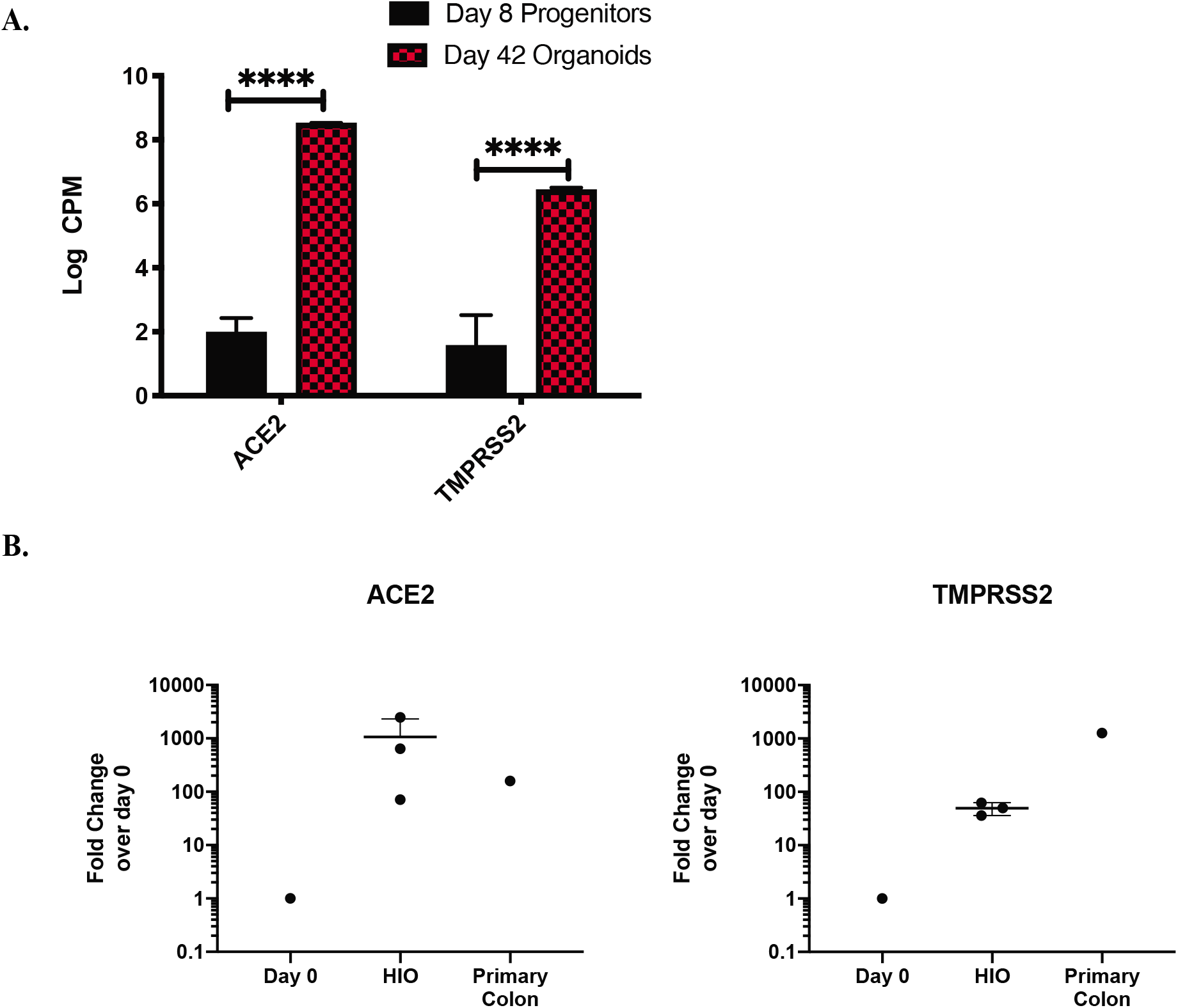
iPSC-derived Human Intestinal Organoids (HIOs) express *ACE2* and *TMPRSS2*. (A) Log Counts Per Million (Log CPM) for *ACE2* and *TMPRSS2* of day 42 HIO compared to day 8 progenitors (n=2 replicates per condition, ****= p<0.0001 calculated as described (Law 2014). HIOs significantly upregulate *ACE2* and *TMPRSS2* compared to day 8 progenitors. (B) qRT-PCR of *ACE2* and *TMPRSS2* in undifferentiated day 0 iPSCs (n=1), day 30 HIOs (n=3 differentiations), and primary human adult colon (n=1). Fold change calculated as 2^-ΔΔCt^ over day 0 sample, error bars represent the s.d.

## Discussion

The unique clinical syndrome COVID-19 with its high rate of ARDS and impressive inflammatory reaction requires urgent attention. Our results indicate that iPSC-derived airway and alveolar epithelial cells cultured at air-liquid interface may serve as a more physiologically relevant model compared to currently available cell lines to study SARS-CoV-2 infection. We found lung cancer cell lines such as Calu-3, A549 and H1299, which have been previously used to model coronavirus infections (Hoffmann 2020, Letko 2020, Matsuyama 2020, Zhang Science 2020), do not express alveolar epithelial-specific transcriptomic programs and may not be sufficient to understand the pathogenesis of SARS-CoV-2 lung infection (Mason 1980). Although we have not yet investigated airway epithelial cell type specific transcriptomic program expression in Calu-3 and A549 cells, the lack of *NKX2-1* expression in these frequently-used lines suggests that they are unlikely to transcriptomically resemble airway epithelial cells. Vero cells, an African Green Monkey kidney cell line, have also been used to model SARS-CoV-2 infection (Hoffmann 2020, Walls 2020, Zhou 2020). Vero cells are readily infectible by SARS-CoV-2 (Hoffmann 2020, Walls 2020, Zhou 2020) and can therefore serve as a model system to study viral entry. However, Vero cells have been described as defective in their ability to mount a type I interferon response to viral infection (Desmyter 1968, Osada 2014) and therefore their utility in understanding the cellular response to SARS-CoV-2 may be limited. Unlike lung cancer cell lines, iPSC-derived airway and alveolar epithelial cells that are cultured at ALI resemble their primary human counterparts transcriptomically and therefore might serve as more physiologically relevant model systems for viral infection.

Importantly, we found that iAT2 and iBC-derived ALI cultures express *ACE2* and *TMPRSS2*, two host factors that permit SARS-CoV-2 cell entry (Hoffmann 2020). Although *ACE2* is expressed in only a minor subset of these cells, expression frequencies in iPSC-derived cell types were commensurate with rates of expression in primary adult human lung cells (Ziegler 2020, Lukassen 2020, Zhang AJRCCM 2020). It is possible that *ACE2* expression is upregulated by interferon induction (Ziegler 2020), and it is not known whether *ACE2* and *TMPRSS2* need to be expressed in the same cell (Courtard 2020, Letko 2020). Additionally, it remains unclear whether SARS-CoV-2 can infect cells independent of ACE2 expression; therefore, further investigations of the mechanisms of SARS-CoV-2 cellular entry is required.

Extrapulmonary cell types may also be susceptible to SARS-CoV-2 infection. Clinically, gastrointestinal manifestations of disease have been observed in COVID-19 patients. In one cohort of 140 patients in Wuhan, China, 40% of COVID-19 patients reported gastrointestinal symptoms (Zhang Allergy 2020). The first confirmed COVID-19 patient in the United States presented with primarily gastrointestinal symptoms and had both fecal and respiratory specimens test positive for SARS-CoV-2 RNA (Holshue 2020). Enterocytes of the small intestine express ACE2 protein (Hamming 2004), and *TMPRSS2* is also highly expressed in human colonic epithelium (Vaarala 2001). SARS-CoV-2 has been identified by electron microscopy in intestinal epithelial cells (Bradley 2020) and viral RNA has been isolated from fecal samples from COVID-19 patients (Xiao 2020, Wang 2020, Holshue 2020). Potential fecal-oral transmission represents an important area of further investigation with significant public health consequences. This extends the necessity of modeling the infection of multiple clinically relevant and molecularly susceptible cell types to include the gastrointestinal epithelium. iPSC-derived intestinal organoids transcriptomically resemble primary gastrointestinal epithelium (Mithal 2020) and express *ACE2* and *TMPRSS2*, suggesting the possibility that iPSC-derived intestinal organoids could also be used to model SARS-CoV-2 infection.

Despite the potential advantages of iPSC-based systems discussed above, we also recognize the limitations. Directed differentiation of human iPSCs to alveolar or airway epithelial-like cells is a time-consuming process (usually more than 40 days) and is technically challenging to master in a short period of time. This is especially inconvenient given the urgency to study SARS-CoV-2 in the midst of the current pandemic. To address this issue, we have developed approaches to facilitate sharing of iAT2s and iBCs as live cell cultures or frozen vials with other researchers. This information is listed under the Resource Sharing section of this manuscript. iAT2s and iBCs can be thawed and expanded in 3D organoid culture and are simple to maintain as spheres or culture at air-liquid interface for experimentation.

In summary, we identify iPSC-derived alveolar and airway epithelial-like cells as a physiologically relevant model system with the potential to model components of SARS-CoV-2 infection such as viral entry, cellular response to pathogen, and viral replication. iAT2s and iBC-derived ALI cultures express the relevant viral entry genes and are an immediately sharable resource. We believe that iAT2s and iBCs can be used to provide insight into lung epithelial cell infection by SARS-CoV-2 and test therapeutic candidates.

### Resource Sharing

#### iAT2 and iBC availability

To request iAT2s or iBCs that can be expanded and maintained at the self-renewing sphere stage of differentiation and to learn more information about our data, differentiation and maintenance protocols for iAT2 and iBCs, please visit http://www.bumc.bu.edu/kottonlab/covid-19-research/.

#### iPSC availability

To view our complete iPSC repository, please visit https://stemcellbank.bu.edu.

#### Data availability

The dataset supporting the conclusions of these experiments is available in the GEO repository, accessions GSE128922, GSE135893 and in the ArrayExpress repository E-MTAB-2706.

## Methods

### RNA-Seq Comparison to Existing Human Lung Cell Line Models

RNA-Seq data from 150 lung cell lines were downloaded from the ArrayExpress repository with accession E-MTAB-2706 (https://www.ebi.ac.uk/arrayexpress/experiments/E-MTAB-2706/) characterized by Klijn et al. (2015). For the comparison with our bulk sequenced primary and iPS derived cells, RPKM (reads per kilobase of transcript per million reads mapped) data were used. The RPKM transformation accounts for differences in both library size and transcript size. RPKM values were log-transformed for visualization purposes. Log-transformation included an offset of 1, to avoid infinite values when taking the logarithm of 0.

### iAT2 Air-Liquid Interface (ALI) culture

iAT2s in 3D spheres were generated as previously described (Jacob 2019) and plated in Matrigel (Corning) droplets at a density of 400 cells/ul. Matrigel droplets were dissolved in 2 mg/ml dispase (Sigma) and alveolospheres were dissociated in 0.05% trypsin (Gibco) to generate a single-cell suspension. 6.5mm Transwell inserts (Corning) were coated with dilute Matrigel (Corning) according to the manufacturer’s instructions. Single-cell iAT2s were plated on Transwells at a density of 520,000 live cells/cm^2^ in 100ul of CK+DCI+Y (3uM CHIR99021, 10ng/ml KGF, 50nM dexamethasone, 0.1mM cAMP, 0.1mM IBMX, 10uM Rho-associated kinase inhibitor (Sigma Y-27632)). 600ul of CK+DCI+Y was added to the basolateral compartment. 24 hours after plating, basolateral media was refreshed to CK+DCI+Y. 48 hours after plating, apical media was aspirated to initiate air-liquid interface culture. 72 hours after plating, basolateral media was changed to CK+DCI to remove the Rho-associated kinase inhibitor. Basolateral media was changed 3 times per week thereafter.

### Intracellular Flow Cytometry for NKX2-1

iAT2 ALIs were dissociated using Accutase (Sigma A6964) according to the manufacturer’s instructions. Cells were evaluated for intracellular NKX2-1 protein by flow cytometry as previously described (Hawkins 2017).

### iAT2 ALI Immunofluorescent Staining

iAT2 ALI cultures were fixed with 4% paraformaldehyde (Electron Microscopy Sciences 19208). Cultures were washed, permeabilized using 0.25% Triton X-100 (Sigma 9002-93-1) for 30 minutes at room temperature, and then blocked with 2.5% normal donkey serum (Sigma D9663) for a total of 1 hour at room temperature. Cells were incubated with anti-pro-SFTPC (Santa Cruz sc-518029, 1:500) primary antibody overnight at 4°C. Cells were washed and incubated with secondary antibody conjugated to Alexa Fluor 488 (Jackson Immunoresearch 711-545-150, 1:500) for 2 hours at room temperature. Nuclei were counterstained with Hoechst 33342 dye (ThermoFisher H3570, 1:500). Transwell membranes were excised with a razor blade and mounted on glass slides using Prolong Diamond Anti-Fade Mounting Reagent (ThermoFisher P36931) and visualized with a Zeiss confocal microscope.

### iAT2 scRNA-Seq

iAT2s were generated from previously published iPSC line (SPC2-ST-B2 also known as “SPC2”; Hurley 2020). 3D iAT2s were dissociated and re-plated either on Transwells for ALI culture or in Matrigel droplets for continued 3D alveolosphere culture as previously described. 10 days after re-plating, passage-matched 3D and ALI iAT2s were dissociated to single cells, live cells were selected using calcein blue and sorted on a Mo-Flo Astrios Cell sorter. Capture and library preparation was performed using a 10X Chromium system. Libraries were sequenced using a NextSeq 500. Where indicated in the text and figures, previously published datasets were also employed (Jacob 2017, Hurley 2020), profiling iAT2s derived from the RUES2 human embryonic stem cells line.

### iBC ALI Culture

iBCs were generated as previously described (Hawkins et al. 2020). We previously generated a basal cell reporter iPSC line, NKX2-1^GFP^;TP63^tdTomato^ (“BU3 NGPT”). Briefly, NKX2-1^GFP^+/TP63^tdTomato^+ iBCs generated from this iPSC line were sorted and plated into air-liquid interface 2D cultures for at least 2 weeks in Pneumacult ALI media where they formed a pseudostratified epithelium composed of multiciliated, secretory and basal cells.

### iBC ALI scRNA-Seq

iBCs from BU3 NGPT cells were generated as above and plated into ALI as previously described (Hawkins 2020). Cells were dissociated to single cells using 0.05% trypsin, live cells selected using calcein blue and sorted on a Mo-Flo Astrios Cell sorter, and captured and library preparation was performed using a 10X Chromium system. Libraries were then sequenced using a NextSeq 500.

### Bioinformatic analysis of the combined data sets

Single cell reads from iAT2 and airway cultures were aligned to the GRCh38 reference genome to obtain a gene-to-cell count matrix using Cell Ranger version 3.0.2 (10x Genomics). The data from Habermann et al. was imported directly as a Seurat object with its annotations; samples from IPF or other diagnosis were filtered out and the remaining healthy controls were then merged with our iAT2 and airway data to form a single matrix.

This matrix was pre-processed and filtered to remove stressed or dead cells and potential doublets based on the percentage of reads mapping to mitochondrial genes (which is a proxy for degradation) and in the number of genes and transcripts detected overall (doublets tend to be outliers for these latter metrics). Normalization, scaling and regressing out unwanted sources of variation (like cell degradation or cell cycle), was done with the SCTransform method (regularized-negative binomial modeling). Subsequently, linear dimensionality reduction (PCA) was performed using the 3000 genes with highest variance. The principal components were then used for batch correction, using the Harmony package to control for variability associated with the data source factor. Clustering was done using the Louvain method for modularity optimization. Further non-linear dimensionality reduction was performed using UMAP for visualization purposes. Clusters identified with the Louvain method were annotated based on their enrichment score for molecular signatures taken from the top 20 differentially expressed genes (computed using hurdle models for sparse single cell data, as implemented in the MAST package) in the dataset from Habermann et al.

### Generation of HIOs, digital gene expression, and qRT-PCR

The iPSC line BU1CG was differentiated to human intestinal organoids (HIOs) and profiled by digital gene expression as previously described (Mithal 2020). qRT-PCR was performed using Taqman probes *ACE2* (ThermoFisher Hs01085333_m1) and *TMPRSS2* (ThermoFisher Hs01122322_m1). RNA from adult human colonic epithelium was obtained from ThermoFisher (#AM7986).

## Acknowledgements

The authors wish to thank all members of the Wilson, Hawkins, Kotton, Mostoslavsky, and Ikonomou Labs for helpful discussions. KMA is supported by NIH grant F30HL147426. RBW is supported by a CJ Martin Early Career Fellowship from the Australian National Health and Medical Research Council. KDA is supported by the I.M. Rosenzweig Junior Investigator Award from the Pulmonary Fibrosis Foundation. AM is supported by NIH Grant 5T32HL007969-12 and the Kilachand Fellows Program at Boston University. GM is supported by NIH Grant 1R24HL123828. FH is supported by Cystic Fibrosis Foundation (CFF) grant HAWKIN19XX0 and NIH grants U01HL148692 and R01HL139799. DNK is supported by an Evergrande MassCPR award, and NIH grants U01HL148692, U01HL134745, U01HL134766, and R01HL095993. AAW is supported by NIH grants U01TR001810, R01DK101501, and R01DK117940. iPSC distribution and disease modeling are supported by NIH grants U01TR001810, and N01 75N92020C00005.

---

**Figure S1.**
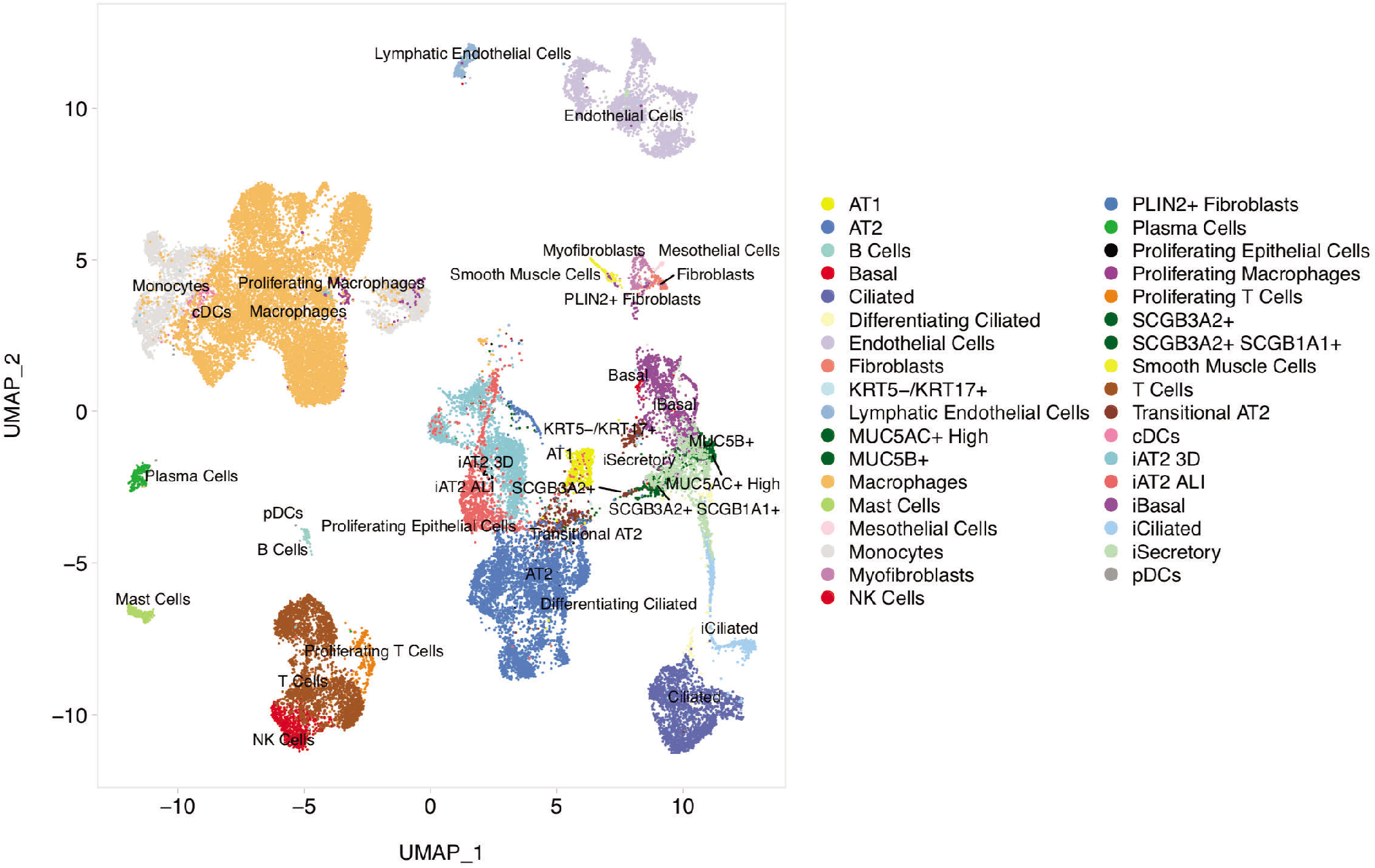
iAT2s and iBC-derived airway epithelial cells are transcriptomically similar to primary counterparts. UMAP with annotations by cell type. The annotations of the primary human lung cells are taken from their original analysis (Habermann 2020). The annotations for the iPSC-derived samples are manually curated based on the expression of known markers and signature enrichments.

